# ASK1 promotes inflammation in senescence and aging

**DOI:** 10.1101/2023.09.20.558729

**Authors:** Takeru Odawara, Shota Yamauchi, Hidenori Ichijo

**Author notes:** Corresponding author. (S.Y.); (H.I.).

## Abstract

Cellular senescence is a stress-induced, permanent cell cycle arrest involved in tumor suppression and aging. Senescent cells secrete bioactive molecules such as pro-inflammatory cytokines and chemokines. This senescence-associated secretory phenotype (SASP) has been implicated in immune-mediated elimination of senescent cells and age-associated chronic inflammation. However, the mechanisms regulating the SASP are incompletely understood. Here, we show that the stress-responsive kinase ASK1 promotes inflammation in senescence and aging. ASK1 is activated during senescence and increases the expression of pro-inflammatory cytokines and chemokines by activating p38, a kinase critical for the SASP. ASK1-deficient mice show impaired elimination of oncogene-induced senescent cells and an increased rate of tumorigenesis. Furthermore, ASK1 deficiency prevents age-associated p38 activation and inflammation and attenuates glomerulosclerosis. Our results suggest that ASK1 is a driver of the SASP and age-associated chronic inflammation and represents a potential therapeutic target for age-related diseases.

## Introduction

Cellular senescence is a state of persistent cell cycle arrest that is induced by various stresses, such as telomere shortening, oncogene activation, and DNA damage^1^. Senescence has long been known as a tumor suppressive mechanism that prevents the proliferation of potentially tumorigenic cells^1^. In addition, recent studies have shown that senescence occurs in various tissues over time and contributes to aging^2^.

Senescent cells exert a variety of physiological functions through the secretion of a complex combination of pro-inflammatory cytokines, chemokines, and growth factors, termed the senescence-associated secretory phenotype (SASP)^3^. The SASP can be beneficial or detrimental depending on the biological context. For example, the SASP is beneficial in that it mediates the recruitment of immune cells, such as macrophages, to eliminate senescent cells^4^. This elimination is called senescence surveillance and is important for tumor suppression^4^. On the other hand, the SASP is detrimental through the contribution to age-associated chronic inflammation^5^. Clearance of senescent cells from tissues reduces the expression levels of pro-inflammatory cytokines and extends lifespan in naturally or accelerated aging mice^6,7^. Conversely, transplantation of senescent preadipocytes into young mice leads to physical dysfunction and shortened lifespan^8^.

Chronic inflammation is a hallmark of aging^9^. Elevated levels of inflammatory cytokines in the elderly are associated with the risk of age-related diseases such as atherosclerosis, diabetes, and neurodegenerative disorders^10–12^. Blockade of TNF-α or knockout of the inflammasome protein NLRP3 suppresses inflammation in aged mice and ameliorates age-related pathologies such as cardiac hypertrophy and decline in motor and cognitive function^13–16^. It has been reported that approximately 40% of the proteins that are increased in aging plasma are associated with the SASP^17^. This implies the potential of drugs that suppress the SASP, called senomorphics, as a treatment for age-related diseases^18^. Therefore, it is important to understand the molecular mechanism of the SASP.

Previous studies have identified several SASP drivers such as NF-κB, C/EBP-β and p38 mitogen-activated protein kinase (MAPK)^19–21^. p38 promotes SASP factor transcription and, at the post-transcriptional level, stabilizes SASP factor mRNAs^21,22^. The importance of p38 in the SASP is supported by the finding that inhibition of p38 reduces the tumor-promoting activities of senescent fibroblasts in the tumor microenvironment^22^. However, the mechanism of p38 activation during senescence is largely unknown.

In this study, we report that the stress-responsive kinase ASK1 activates p38 during senescence and potentiates the expression of SASP factors, thereby mediating the tumor suppressor function of the SASP. Moreover, we reveal that the ASK1-p38 pathway is also activated during aging. In aged mice, ASK1 increases the expression of IL-1β and promotes glomerulosclerosis. Our findings demonstrate that ASK1 activation is crucial for inflammation associated with senescence and aging.

## Results

### ASK1 promotes the SASP by activating p38

In the MAPK cascade, MAP3K phosphorylates and activates MAP2K, which in turn phosphorylates and activates MAPK. To identify the activation mechanism of p38 in cellular senescence, we performed an siRNA screen for MAP3Ks involved in the SASP. We used IMR-90 primary human fibroblasts in which KRAS^G12V^ expression is inducible with 4-hydroxy Tamoxifen (4OHT) (hereafter IMR-90 ER-KRAS), a well-characterized model of oncogene-induced senescence (OIS)^23^. We also induced senescence by treating IMR-90 cells with the DNA-damaging agent doxorubicin. Of the MAP3Ks expressed in IMR-90 cells, knockdown of MAP3K5 (ASK1) suppressed senescence-associated upregulation of IL8 and IL1B in both OIS and DNA damage-induced senescence (DIS) (Fig. 1a).

**Fig. 1.**
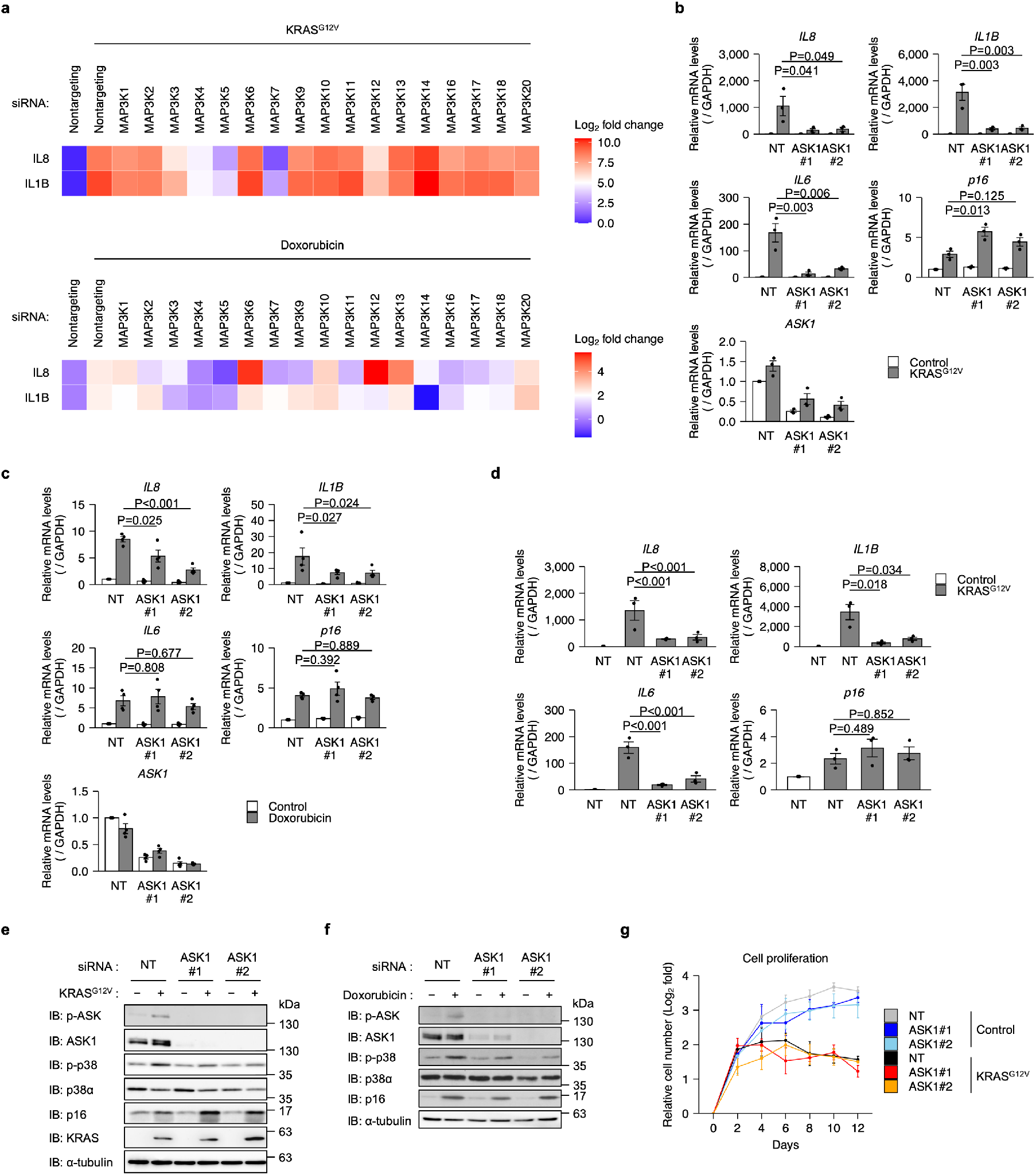
ASK1 promotes the SASP by activating p38. **a**, Heatmap of the mRNA levels for IL8 and IL1B. Cells were transfected with non-targeting (NT) or indicated MAP3K siRNAs and treated with 4OHT (IMR-90 ER-KRAS) or doxorubicin (IMR-90) for 12 days. The mRNA levels were analyzed by qPCR. *n* = 3 (top), 2 (bottom). Data are mean. **b**, qPCR analysis of IMR-90 ER-KRAS cells transfected with NT and ASK1 siRNAs and treated with 4OHT for 12 days. *n* = 3. **c**, qPCR analysis of IMR-90 cells transfected with NT and ASK1 siRNAs and treated with doxorubicin for 12 days. *n* = 4. **d**, qPCR analysis of IMR-90 ER-KRAS cells transfected with NT and ASK1 siRNAs at day 8 after 4OHT treatment. *n* = 3. Bars represent mean ± s.e.m. (**b-d**). **e, f**, Immunoblot analysis of indicated cells transfected with NT and ASK1 siRNAs and treated with 4OHT (**e**) or doxorubicin (**f**) for 12 days. **g**, Growth curves of IMR-90 ER-KRAS cells transfected with NT and ASK1 siRNAs and treated with 4OHT or DMSO. *N* = 3. Data are mean ± s.e.m. Statistical analysis was performed using Dunnett’s multiple comparison test (**b-d**).

ASK1 is a MAP3K upstream of p38 and JNK and has been implicated in various stress responses such as cytokine production in innate immunity^24^. We confirmed by qPCR that knockdown of ASK1 suppressed expression of SASP factors in both OIS and DIS (Fig. 1b, c). Knockdown of ASK1 in already senescent cells also decreased expression of SASP factors (Fig. 1d). Immunoblotting analysis revealed that ASK1 and p38 were activated during OIS and DIS (Fig. 1e, f). p38 activation was suppressed by knockdown of ASK1 (Fig. 1e, f). In contrast, increased expression of p16, a cyclin-dependent kinase inhibitor critical for senescence, was not suppressed by ASK1 knockdown at either the mRNA or protein level (Fig. 1b-f). Knockdown of ASK1 did not significantly affect cell proliferation in 4OHT-treated IMR-90 ER-KRAS cells (Fig. 1g), suggesting that ASK1 is dispensable for the senescence proliferative arrest. These data indicate that ASK1 promotes the SASP via activation of p38.

### ASK1 mediates macrophage migration through the SASP

We next sought to explore the role of ASK1 in biological functions of the SASP. Senescent cells attract macrophages and other immune cells through the SASP, thereby promoting their own elimination^4,25,26^. To examine the involvement of ASK1 in macrophage recruitment, we performed a cell migration assay using THP-1 cells (Fig. 2a). Conditioned medium (CM) derived from senescent IMR-90 ER-KRAS cells promoted the migration of THP-1 cells compared to CM derived from proliferative cells (Fig. 2b). This migration was inhibited by knockdown of ASK1 in senescent IMR-90 ER-KRAS cells, suggesting that ASK1 is required for the attraction of macrophage through the SASP *in vitro* (Fig. 2b).

**Fig. 2.**
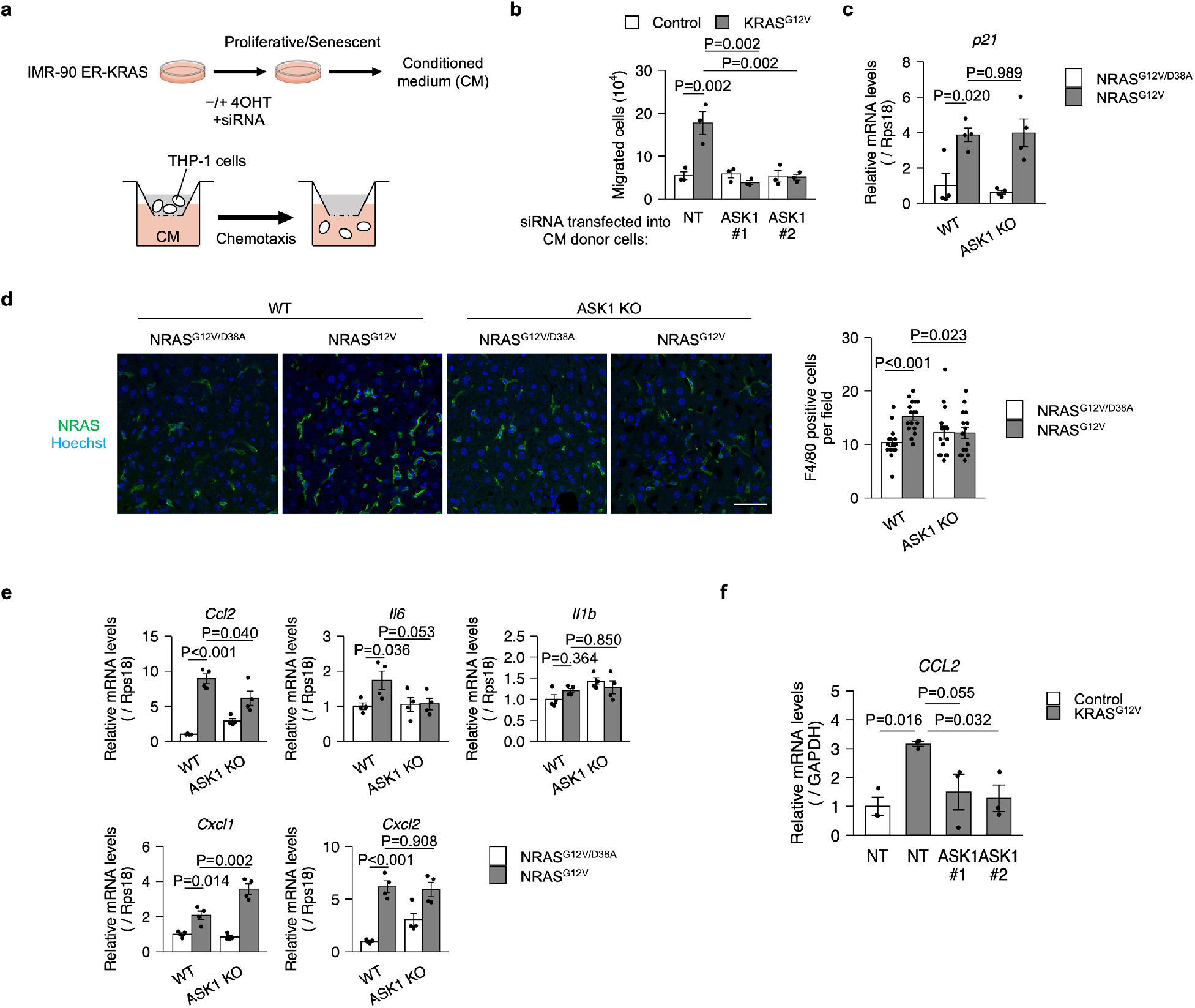
ASK1 mediates macrophage migration through the SASP. **a**, Schematic of the migration assay of THP-1 cells with proliferative and senescent IMR-90 ER-KRAS. CM, conditioned media. **b**, Migration assay of THP-1 cells. CM was collected from proliferative and senescent IMR-90 ER-KRAS cells transfected with non-targeting (NT) and ASK1 siRNAs. *n* = 3. **c, d**, qPCR analysis (**c**) and representative immunofluorescence images (**d**) of wild-type (WT) and ASK1-knockout (ASK1 KO) mice liver 6 days after hydrodynamic gene delivery. *n* = 4 (**c**), 3 (**d**) mice per group. 5 randomly selected fields per mouse were analyzed. Scale bar, 50 μm. **e**, qPCR analysis of WT and ASK1 KO mice liver 6 days after hydrodynamic gene delivery. *n* = 4 mice per group. **f**, qPCR analysis of IMR-90 ER-KRAS cells transfected with NT and ASK1 siRNAs and treated with 4OHT for 12 days. *n* = 3. Bars represent mean ± s.e.m. (**b-f**). Statistical analysis was performed using Dunnett’s multiple comparison test (**b, c, e, f**) and unpaired Student’s t-test (**d**).

To investigate whether ASK1 is involved in macrophage recruitment *in vivo*, we used a mouse model in which hepatocyte senescence is induced by hydrodynamic injection-mediated delivery of an oncogenic NRAS vector (NRAS^G12V^)^4,27,28^. Consistent with previous studies, expression of NRAS^G12V^, but not an inactive mutant of NRAS (NRAS^G12V/D38A^), upregulated the expression of the senescence marker p21 at day 6 (Fig. 2c). p21 expression levels were comparable between wild-type (WT) and ASK1-knockout (ASK1 KO) mice, indicating that ASK1 does not affect senescence induction (Fig. 2c). In contrast, immunostaining of liver sections revealed that macrophage recruitment was significantly impaired in ASK1 KO mice (Fig. 2d). In this experimental model, increased CCL2 expression in NRAS^G12V^-expressing hepatocytes was shown to promote macrophage recruitment^4,27,28^. This increased CCL2 expression was suppressed in ASK1 KO mice (Fig. 2e). In addition, increased CCL2 expression in senescent IMR90 ER-KRAS cells was suppressed by knockdown of ASK1 (Fig. 2f). These data indicate that ASK1 is involved in the SASP-mediated recruitment of macrophages.

### ASK1 is required for the tumor suppressor function of the SASP

Impaired macrophage recruitment results in incomplete elimination of senescent cells^4^. Using a luciferase reporter co-expressed with the NRAS mutants, we investigated the requirement of ASK1 for the elimination of senescent hepatocytes. WT mice showed NRAS activity-dependent reduction in luciferase activity from day 6 to day 12 (Fig. 3a). This reduction was partially but significantly inhibited in ASK1 KO mice (Fig. 3a). Re-expression of ASK1 along with NRAS^G12V^ in hepatocytes of ASK1 KO mice restored the reduction in luciferase activity (Fig. 3b). OIS cells that evade senescence surveillance can develop into tumors^4^. Five months after injection of the NRAS^G12V^ vector, more intrahepatic tumors were observed in ASK1 KO mice than in WT mice (Fig. 3c). These data demonstrate that ASK1 is crucial for the elimination of pre-cancerous senescent cells.

**Fig. 3.**
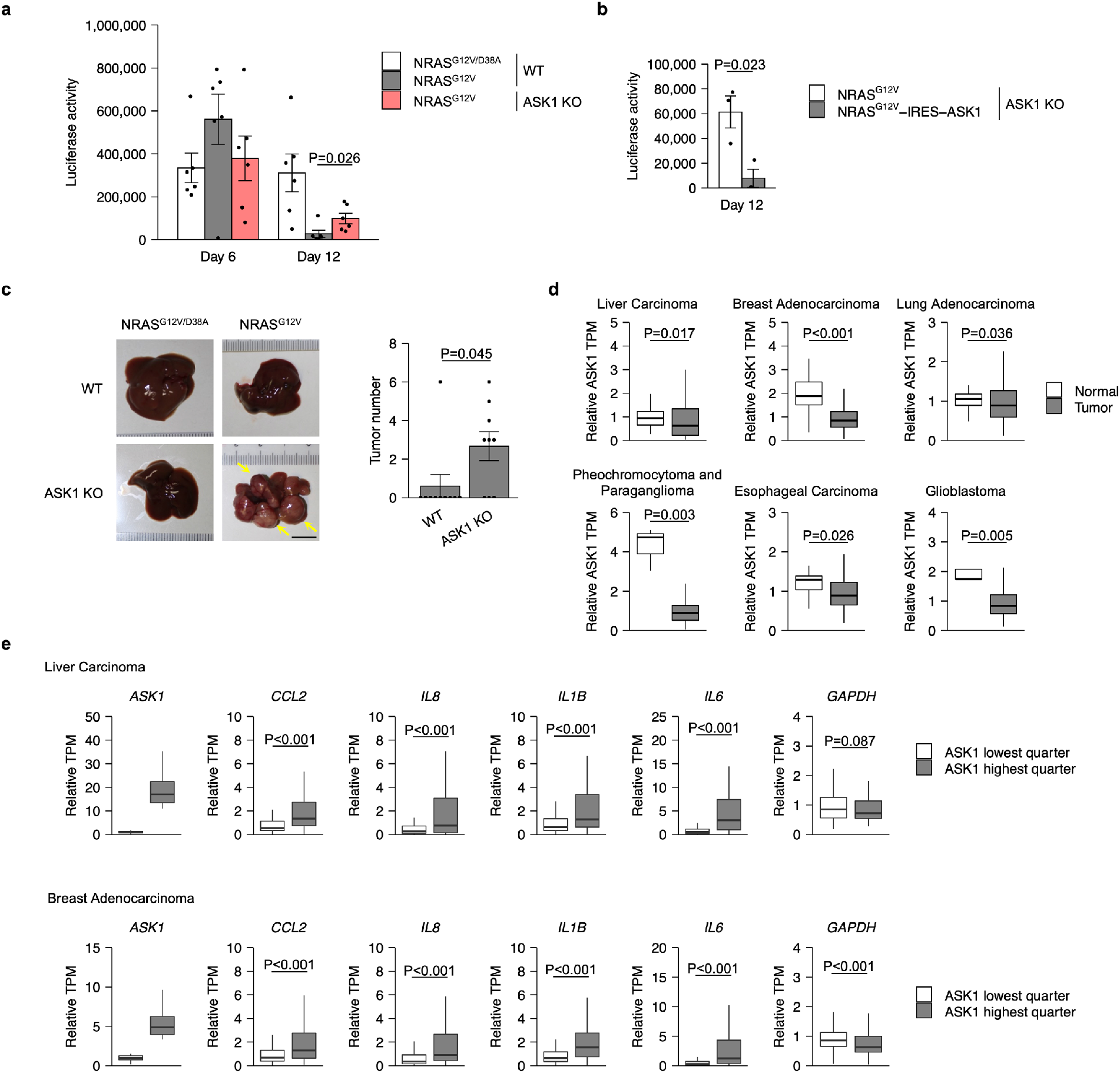
ASK1 is required for the tumor suppressor function of the SASP. **a**, Luciferase assay of wild-type (WT) and ASK1-knockout (ASK1 KO) mice liver after hydrodynamic gene delivery. *n* = 6 mice per group. **b**, Luciferase assay of ASK1 KO mice liver 12 days after hydrodynamic gene delivery. *n* = 3 mice per group. **c**, Representative images of WT and ASK1 KO mice liver 5 months after hydrodynamic gene delivery. *n* = 9 (WT), 10 (ASK1 KO) mice per group. Yellow arrows indicate tumors. Scale bar, 1 cm. Bars represent mean ± s.e.m. (**a-c**). **d**, RNA-seq analysis of ASK1 expression in normal and tumor tissues. RNA-Seq data were obtained from the TCGA project. **e**, RNA-Seq analysis of inflammatory gene expression in tumor tissues with high and low ASK1 expression. RNA-seq data from indicated TCGA project were used for the analysis. Statistical analysis was performed using Wilcoxon rank-sum test (**a, c, d, e**) and unpaired Student’s t-test (**b**).

To investigate the role of ASK1 in human cancer, we explored the RNA-seq data in The Cancer Genome Atlas (TCGA). In several cancer types, ASK1 expression was lower in cancer tissues compared to normal tissues (Fig. 3d). In liver and breast cancer, low expression of ASK1 was associated with low expression of pro-inflammatory cytokines and chemokines (Fig. 3e). These data are consistent with the idea that, at least in certain human cancers, ASK1 deficiency impairs senescence surveillance and promotes tumorigenesis.

### ASK1-p38 pathway promotes age-related inflammation

The SASP is associated with chronic inflammation and diseases during aging^11^. In particular, previous reports indicate that an age-associated increase in serum concentrations of IL-1β contributes to diseases such as type 2 diabetes, Alzheimer’s disease, and motor dysfunction^16,29,30^. In ASK1 KO mice, increased IL-1β expression with age tended to be suppressed in the kidney, lung, brain, and muscle (Fig. 4a). In contrast, p16 expression increased with age in both WT and ASK1 KO mice, except in muscle (Fig. 4b). ASK1 and p38 were activated in the kidney and lung of aged mice (Fig. 4c, d). Furthermore, p38 activation was suppressed in ASK1 KO mice (Fig. 4c, d). Although ASK1 activity could not be evaluated in the brain, age-associated p38 activation was suppressed in ASK1 KO mice (Fig. 4c). These data indicate that the ASK1-p38 pathway is activated during aging and contributes to pro-inflammatory cytokine production.

**Fig. 4.**
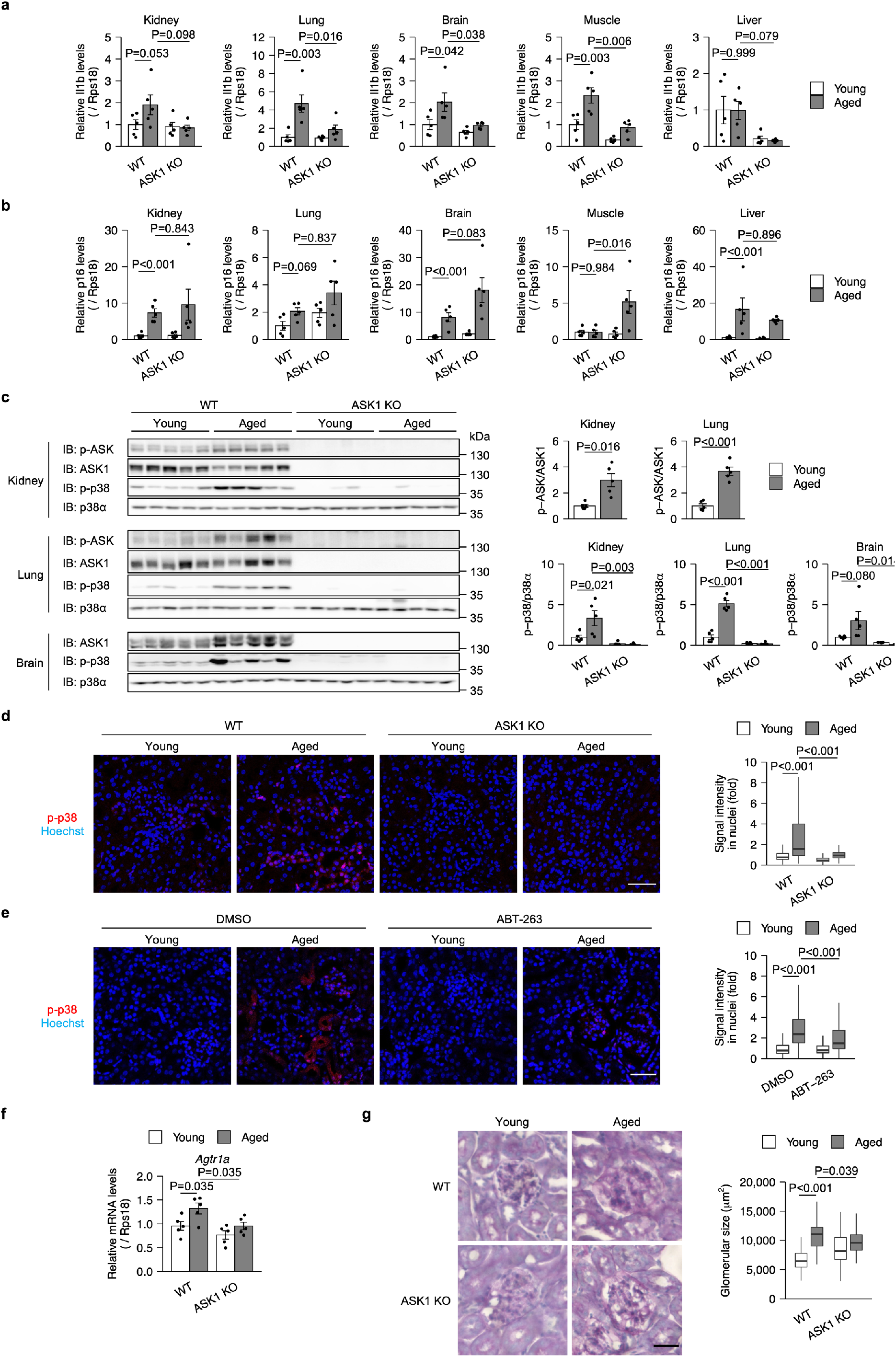
ASK1 is involved in age-related inflammation. **a, b**, qPCR analysis of *Il1b* (a) and *p16* (b) in the indicated tissues from young (3-month-old) and aged (20-month-old), wild-type (WT) and ASK1-knockout (ASK1 KO) mice. *n*= 5 mice per group. **c**, Immunoblot analysis (left) and band quantification (right) of the indicated tissues from WT and ASK1 KO mice. *n* = 5 mice per group. **d**, Representative immunofluorescence images (left) and quantification (right) of kidneys from WT and ASK1 KO mice. Scale bar, 50 μm. *n* = 3 mice per group. 10 randomly selected fields per mouse were analyzed. **e**, Representative immunofluorescence images (left) and quantification of phospho-p38 signal (right) of kidneys from young and aged WT mice treated with DMSO or ABT-263. 10 randomly selected fields per mouse were analyzed. Scale bar, 50 μm. *n* = 3 mice per group. **f**, qPCR analysis of young and aged mouse kidneys from young and aged, WT and ASK1 KO mice. *n* = 5 mice per group. **g**, Representative images of periodic acid-Schiff (PAS) staining (left) and quantification of glomerular size (right) of kidneys from young and aged, WT and ASK1 KO mice. Bars represent mean ± s.e.m. (**a-f**). Scale bar, 50 μm. *n* = 3 mice per group. Statistical analysis was performed using Dunnett’s multiple comparison test (**a-g**).

Senescent cells that accumulate in aged kidney tissue cause age-associated renal inflammation and dysfunction^7^. We asked whether cellular senescence is required for the activation of the ASK1-p38 pathway in aged kidney. To this end, we induced selective elimination of senescent cells, termed senolysis. Immunohistochemical analysis revealed that peritoneal injection of ABT-263, a widely used senolytic drug, reduced phospho-p38 signal in aged kidney (Fig. 4e), suggesting that senescent cells are involved in the activation of the ASK1-p38 pathway during renal aging.

Glomerulosclerosis is a hallmark of age-related renal inflammation^31,32^. As a possible cause of glomerulosclerosis, it has been reported that senescent cells increase the expression of the angiotensin receptor Agtr1a in the kidney^7^. We confirmed that Agtr1a expression increased with age (Fig. 4f). This increase was impaired in ASK1 KO mice (Fig. 4f). Furthermore, the glomerular enlargement characteristic of glomerulosclerosis was ameliorated in ASK1 KO mice^33^ (Fig. 4g). These data indicate that ASK1 plays a key role in age-related renal inflammation.

## Discussion

The SASP has been suggested as a contributor to age-associated inflammation and a potential therapeutic target for age-related diseases^18^. Previous studies have shown that pharmacological inhibition of NF-κB and other SASP drivers ameliorates ataxia, kidney glomerulosclerosis, osteoporosis, and frailty^34–37^. However, until recently, there was no genetic evidence that the molecular mechanisms that regulate the SASP are also involved in age-associated inflammation^38^. In the present study, we found that ASK1 is required for both the SASP and age-associated inflammation.

Using pathological mouse models, we have reported that ASK1 is involved in age-related diseases such as cardiac hypertrophy and Parkinson’s disease^39,40^. Furthermore, p38 has been implicated in age-related diseases. In muscle stem cells, p38 is activated with age and decreases their proliferative capacity^41,42^. p38 is also activated in aged and Alzheimer’s disease brains^43,44^. Therefore, chronic inflammation induced by the ASK1-p38 pathway may contribute to a variety of age-related diseases. ASK1 inhibition has been found to be well tolerated in clinical trials and may be a promising future therapeutic option for age-related diseases^45^.

## Methods

### Cell culture and treatments

IMR-90 normal human fibroblasts (ATCC, CCL-186) and HEK293T cells were cultured in DMEM (Sigma-Aldrich, D5796; Wako, 044-29765) supplemented with 10% FBS (BioWest, S1560-500) and 100 units/ml penicillin G (Meiji Seika). THP-1 human monocytes were cultured in RPMI-1640 (Wako, 189-02025) supplemented with 10% FBS in a 5% CO2 atmosphere at 37 °C. For induction of the OIS, 4-hydroxy Tamoxifen (Cayman Chemical, 17308) at a final concentration of 200 nM was added to IMR-90 ER-KRAS cells. For induction of the DIS, 200 ng/mL doxorubicin hydrochloride (Wako, 040-21521) were added to IMR-90 cells.

### Lentivirus infection

For lentivirus production, HEK293T cells were transfected with the lentiviral vector pLenti-PGK-ER-KRAS^G12V^ (Addgene, 35635) and the packaging plasmids pCMV-VSV-G (Addgene, 8454) and psPAX2 (Addgene, 12260) using PEI MAX transfection reagent (Polyscience, 24765) for 24 h. The medium was replaced with 5 mL of fresh medium containing 1% bovine serum albumin (BSA, Iwai Chemical, A001) and collected 24 h later. This step was repeated. The collected media were combined and filtered through a 0.45 μm PVDF filter (Millipore, SLHVR33RS). IMR-90 cells were infected with lentiviruses in the presence of 8 μg/mL Polybrene (Nacalai Tesque, 17736-44) for 24 h, cultured in fresh media for 24 h, and selected with 100 μg/mL hygromycin (Nacalai Tesque, 07296-11) for 3 days.

### RNA isolation and qPCR

Total RNA was extracted using Isogen (Wako, 319-90211) and subjected to reverse transcription using a ReverTra Ace master mix (Toyobo, FSQ-301). qPCR was performed using a SYBR FAST qPCR kit (KAPA Biosystems, KK4602) and a QuantStudio 1 qPCR system (Applied Biosystems). The following primers were used: *GAPDH*, forward, 5′-AGCCACATCGCTCAGACAC-3′, reverse, 5′-GCCCAATACGACCAAATCC-3′; *ASK1*, forward, 5′-TGAGAAACCTAATGGAATCTTTAGC-3′, reverse, 5′-TGAGGGTTGTGATGTGTTCC-3′; *p16*, forward, 5′-CCAACGCACCGAATAGTTACG-3′, reverse, 5′-GCGCTGCCCATCATCATG-3′; *IL6*, forward, 5′-CCGGGAACGAAAGAGAAGCT-3′, reverse, 5′-GCGCTTGTGGAGAAGGAGTT-3′; *IL8*, forward, 5′-CTTTCCACCCCAAATTTATCAAAG-3′, reverse, 5′-CAGACAGAGCTCTCTTCCATCAGA-3′; *IL1B*, forward, 5′-TACCTGTCCTGCGTGTTGAA-3′, reverse, 5′-TCTTTGGGTAATTTTTGGGATCT-3′; *CCL2*, forward, 5′-AGTCTCTGCCGCCCTTCT-3′, reverse, 5′-GTGACTGGGGCATTGATTG-3′; *mRps18*, forward, 5′-TCCAGCACATTTTGCGAGTA-3′, reverse, 5′-CAGTGATGGCGAAGGCTATT-3′; *mCcl2*, forward, 5′-GTGGGGCGTTAACTGCAT-3′, reverse, 5′-CAGGTCCCTGTCATGCTTCT-3′; *mIl6*, forward, 5′-GCTACCAAACTGGATATAATCAGGA-3′, reverse, 5′-CCAGGTAGCTATGGTACTCCAGAA-3′; *mp21*, forward, 5′-CTGGTGATGTCCGACCTGTT-3′, reverse, 5′-TCAAAGTTCCACCGTTCTCG-3′; *mp16*, forward, 5′-CCCAACGCCCCGAACT-3′, reverse, 5′-GCAGAAGAGCTGCTACGTGAA-3′; *mIl1b*, forward, 5′-AAGAGCTTCAGGCAGGCAGTATCA-3′, reverse, 5′-ATGAGTCACAGAGGATGGGCTCTT-3′; *mCxcl1*, forward,5′-CCCGCTCGCTTCTCTGT-3′, reverse, 5′-CTTTTGGACAATTTTCTGAACCAAG-3′; *mCxcl2*, forward, 5′-CACCAACCACCAGGCTACAG-3′, reverse, 5′-GCTTCAGGGTCAAGGCAAAC-3′; *mAgtr1a*, forward, 5′-AAGGGCCATTTTGCTTTTCT-3′, reverse, 5′-AACTCACAGCAACCCTCCAA-3′. Transcript levels were normalized to *GAPDH* (human) and *mRps18* (mouse).

### siRNA transfection

siRNAs with the following target sequences were purchased from Dharmacon: NT #1 (hereafter ON-TARGETplus), 5′-UGGUUUACAUGUCGACUAA-3′; ASK1 #1 5′-GGGAAUCUAUACUCAAUGA-3′; ASK1 #2 5′-ACACUACAGUCAGGAAUUA-3′.

For the knockdown of MAP3Ks, siRNAs (ON-TARGETplus SMARTpool) targeting each MAP3Ks were purchased from Dharmacon. siRNA transfection was performed at a final concentration of 10 nM using Lipofectamine RNAiMAX transfection reagent (Invitrogen, 13778500) and Opti-MEM (Gibco, 31985070).

### Immunoblotting

Cells were lysed in RIPA buffer (50 mM Tris-HCl [pH 8.0], 150 mM NaCl, 1% NP-40, 0.5% sodium deoxycholate, 0.1% sodium dodecyl sulfate [SDS], 1 mM phenylmethylsulfonyl fluoride, 5 μg/mL leupeptin, 8 mM NaF, 12 mM β-glycerophosphate, 1 mM Na_3_VO_4_, 1.2 mM Na_2_MoO_4_, 5 μM cantharidin, 2 mM imidazole). Lysates were clarified by centrifugation at 17,500 × g for 10 min at 4 °C. The protein concentrations of the lysates were quantified using a BCA protein assay kit (Wako, 297-73101) and equalized. The lysates were then mixed with 2 × SDS sample buffer (125 mM Tris-HCl [pH 6.8], 4% SDS, 20% glycerol, 200 μg/mL bromophenol blue, 10% β-mercaptoethanol) and heated at 98 °C for 3 min. SDS samples were subjected to SDS-polyacrylamide gel electrophoresis (SDS-PAGE) and transferred to Immobilon-P membranes (Millipore, IPVH00010). The membranes were blocked with 5% skim milk (Megmilk Snow Brand) in TBS-T (50 mM Tris-HCl [pH8.0], 150 mM NaCl, 0.05% Tween 20) for 30 min at room temperature and incubated with primary antibodies in TBS-T containing 5% BSA and 0.1% NaN_3_ overnight at 4 °C. The membranes were then incubated with secondary antibodies in TBS-T containing 5% skim milk for 2 h at room temperature, followed by detection using ECL Select detection reagent (Amersham, RPN2235) and a FUSION Solo S chemiluminescence imaging system (Vilber). The following primary antibodies were used: anti-ASK1 (Abcam, ab45178), anti-phospho-p38 (Cell Signaling, #4511), anti-p38α (Cell Signaling, #9228), anti-p16 (Abcam, ab108349), anti-KRAS (Santa Cruz Biotechnology, sc-30), anti-α-tubulin (Bio-Rad, MCA77G). Anti-phospho-ASK antibody was generated as previously described^46^.

### Transwell migration assay

To generate CM, IMR-90 ER-KRAS cells were treated with or without 4OHT for 8 days and then reseeded. The next day, the cells were washed with PBS and then cultured in FBS-free RPMI-1640 for 3 days. Migration assays were performed using transwell polycarbonate membrane inserts with 8 μm pore diameter (FALCON, 353097). 750 μL of CM was added to the bottom chambers. THP-1 cells (5 × 10^5^ cells) resuspended in FBS-free RPMI-1640 medium were added to the inserts. THP-1 cells were allowed to migrate into the bottom chamber for 6 h at 37 °C and the number of cells that had migrated to the bottom chamber was counted.

### Mice

WT and ASK1 KO mice on the C57BL/6 background were described previously^47^. Mice were maintained in a specific pathogen-free facility and age-matched mice were used for the experiment. All the experiments were performed following the experimental protocol approved by the animal ethics committee the University of Tokyo.

### Plasmids

To generate pT/Caggs-NRASV12/D38A, a point mutation was introduced into pT/Caggs-NRASV12 (Addgene, #20205) by PCR using a PrimeSTAR Mutagenesis Basal Kit (Takara, R046A) and the following primers; forward, 5′-ATAGAGGCTTCTTACAGAAAACAAGTG-3′, reverse, 5′-GTAAGAAGCCTCTATGGTGGGATCATA-3′.

### Hydrodynamic gene delivery

Mice aged 8-10 weeks of age were subjected to hydrodynamic injection. 25 μg of pT/Caggs-NRASV12 or pT/Caggs-NRASV12/D38A and 5 μg of PT2/c-Luc//PGK-SB-13 (Addgene, 20207) were suspended in 0.9% saline solution at a final volume of 10% of the body weight and injected via the tail vein within 8 seconds. Vectors were prepared using a QIAGEN EndoFree Mega Kit (QIAGEN).

### Immunohistochemistry

Mouse tissues were perfused and immersed O/N in 4% paraformaldehyde/PBS for fixation, then cleared with 20% sucrose solution. Fixed tissues were embedded in CryoMount I (Muto PureChemicals) and 8 μm-thick sections were cut using a cryostat (Leica). For immunofluorescence assay, sections were blocked in 3% BSA/PBS for 30 min at room temperature, and then incubated with anti-F4/80 (Abcam, ab6640) and anti-phospho-p38 (Cell Signaling, #4511) in 3% BSA/PBS for O/N at 4 °C after antigen retrieval. Antigen retrieval was carried out with sodium citrate buffer (pH 6.0) for 20 min at 92 °C. Then the sections were incubated with Alexa Fluor secondary antibodies (1:100, Invitrogen) and Hoechst 33342 (1:1000) for 2 h at room temperature. Images were acquired using a TCS SP5 confocal microscope (Leica) with 40 × 1.25 NA and 63 × 1.4 NA oil immersion objectives (Leica). Periodic acid-Schiff staining was performed using a PAS staining kit (Sigma-Aldrich, 101646) according to the manufacturer’s instructions. Kidney sections were then counterstained with hematoxylin (Wako, 131-09665).

### Luciferase activity assay

Mouse livers were homogenized in Luciferase Culture Lysis Reagent (Promega). Lysates were clarified by centrifugation at 13,000 × g for 20 min at 4 °C. Supernatants were collected and analyzed using a Luciferase Assay System (Promega). Luminescence was measured using a Varioskan Flash (Thermo Fisher Scientific).

### TCGA data analyses

For TCGA analyses, RNA-seq datasets were obtained from cBioportal (http://www.cbioportal.org). For a given cancer type, the expression level of each gene was normalized to TPM values and ASK1 expression levels were compared between normal and tumor samples. Tumor samples were ranked based on ASK1 expression levels and were evenly divided into four groups. Statistical comparisons were performed between the first group (samples with the lowest 25% expression) and the last group (samples with the highest 25% expression) for inflammatory genes or GAPDH.

### Senolysis

ABT-263 was dissolved in DMSO and then diluted to create a working solution (30% propylene glycol, 5% TWEEN 80, 3.3% dextrose in water, pH 4-5). For the selective elimination of senescent cells in mice, 50 mg/kg body weight of ABT-263 (Chemietek, CT-A263) or DMSO was injected intraperitoneally for 2 consecutive days. Mice were sacrificed 5 days after the last treatment.

### Quantification and statistical analysis

R software was used for statistical analysis. Statistical tests are indicated in the figure legends. P < 0.05 was considered statistically significant. No statistical method was used to predetermine sample size.

## Acknowledgements

We thank M. Fujimoto and M. Kamiyama for assistance with hydrodynamic injection. This work was supported by grants from the Japan Agency for Medical Research and Development (JP21gm5010001 to H.I.), the Japan Science and Technology Agency (JPMJMS2022-18 to H.I.), and the Japan Society for the Promotion of Science (JP18H03995 and JP21H04760 to H.I. and 18K14648 and 21K06061 to S.Y.). T.O. was supported by a SPRING fellowship from the Japan Science and Technology Agency (JPMJSP2108).

## Author contributions

T.O. designed and performed the experiments and analyzed the data. S.Y. and H.I. conceived the study. T.O., S.Y., and H.I. wrote the manuscript.

## Competing interests

The authors declare no competing interests.

**Correspondence and requests for materials** should be addressed to S.Y. or H.I.

## Notes

### Competing Interest Statement

The authors have declared no competing interest.

